# Critical role of DNA unwinding and Cas9 conformational changes in modelling off-target effects

**DOI:** 10.1101/2021.09.28.462201

**Authors:** Aset Khakimzhan, Vincent Noireaux

## Abstract

CRISPR-Cas9 off-target effects interfere with the ability to accurately perform genetic edits. To predict off-target effects CRISPR-Cas9 researchers perform high throughput guide RNA mismatch and bulge experiments and then use the data to fit thermodynamic binding models. While impactful from an engineering perspective such models are not based on the experimentally observed target interrogation process and thus incorrectly measure the energetic effects mismatches have on the system. In this work we convert an experimentally deduced qualitive model of target interrogation to a linear ODE model and demonstrate that the mismatch tolerance patterns observed in experiments do not need to be caused by differences in energetic penalties of mismatches but rather are emergent effects of the timing and coordination of target DNA unwinding and Cas9 conformational changes.

## Introduction

A major bottleneck in the use of CRISPR-Cas9 lies in the off-target effects [1]. Off-target effects are caused by a CRISPR-Cas9 complex binding and cleaving an unintended site on the genome. One strategy to decrease off-target effects is the development of thermodynamic guide RNA optimizing models trained on large mismatch and bulge guide RNA datasets [2–6]. By comparing the cleavage/binding/indel rates between fully complementary guides and mismatched guides researchers inferred the energetic costs accrued by each spacer mutation. While such models are widely used for guide RNA development for Cas9 (and Cas12 [7]) systems they have one major flaw: they ignore much of the latest mechanistic experimental results of CRISPR-Cas9 target interrogation. This approach is understandable for the early stages of CRISPR-Cas9 models but now that the field has acquired a wide range of mechanistic experimental evidence it is important that off-target effect models account for them as well.

In this work we develop a CRISPR-Cas9 model that is based on the latest experimental target interrogation results. An important distinction of this model is the accounting for the discrete steps in which Cas9 unwinds the target DNA. This element of the model is based on structural experiments [8–12], single molecule Forster resonance energy transfer (smFRET) experiments [13–17], molecular dynamics simulations [18–20], and magnetic tweezers experiments [21]. In addition to nucleic acid dynamics the model also accounts for the strategy by which CRISPR-Cas9 searches for the target DNA [22–24]. We demonstrate that the PAM-proximal mismatch tolerance is largely controlled by CRISPR-Cas9 rebinding conditions, while the middle-target and PAM-distal mismatch patterns are dependent of the rates of CRISPR-Cas9 conformational changes. The message of the model is that the mismatch tolerance profiles observed across multiple experiments are emergent of the topology and timing of CRISPR-Cas9’s target recognition process rather than the energies of bonds themselves. In addition to providing a qualitative picture of different mismatch profiles our results also demonstrate how Cas9 conformational changes can be thought of nature leveraging a speed-accuracy tradeoff. With all the parameters of the model explained we test the model against available mismatch data. To reduce the number of parameters, we convert our model into a previously described energy landscape scheme [25] and then utilize this energy landscape to fit data from 30 targets in HEK293T cells [26] using realistic parameter constraints.

### Model of CRISPR-Cas9 target interrogation

The first steps of CRISPR-Cas9 cleavage are target search. Our model relies on the hypothesis that CRISPR-Cas9 search is a mix of 3D diffusion across the larger experimental space and 1D-like sliding between neighboring PAM-sites [23,24,27]. Previous theoretical works studying the dynamics of nucleic acid binding proteins [28] and specifically CRISPR-Cas9 [29] demonstrates that an element of 1D sliding/PAM localization is an important contribution to the search mechanism. The system starts in the overall experimental space (Fig 1A) and via 3D diffusion enters an arbitrary vicinity (Fig 1B) of the target at rate of *k*_*S*_, with the rate of exiting that vicinity to the overall experimental space being k_Lost_. The 1D motion corresponds to the binding and unbinding to the target DNA once in the arbitrary vicinity, labeled *k*_*ON*_ and *k*_*OFF*_ respectively. For our model we set *k*_*S*_ to a realistic for cells value of 10^−2^ s^-1^, which correspond to an average target search time of 100 seconds. The rate of escaping the target vicinity to the general experimental space, *k*_*Lost*_ is set to 1s^-1^, which corresponds to a residence time of 1s. That is a realistic value based on theoretical analysis of CRISPR-Cas9 localization [29]. The measured residence time of non-target is on the order of 30ms [22–24] and therefore the rate of breaking a PAM bond, k_OFF_, can be approximated as 30 s^-1^. The neighboring PAM sites will also have a residence time of approximately 30ms and considering a 1D sliding model a ½ probability of returning to the target site. Therefore an expected timeframe for which CRISPR-Cas9 will rebind to the correct target is on the scale of a 100ms and thus we approximate k_ON_ as 10 s^-1^.

**Figure 1.**
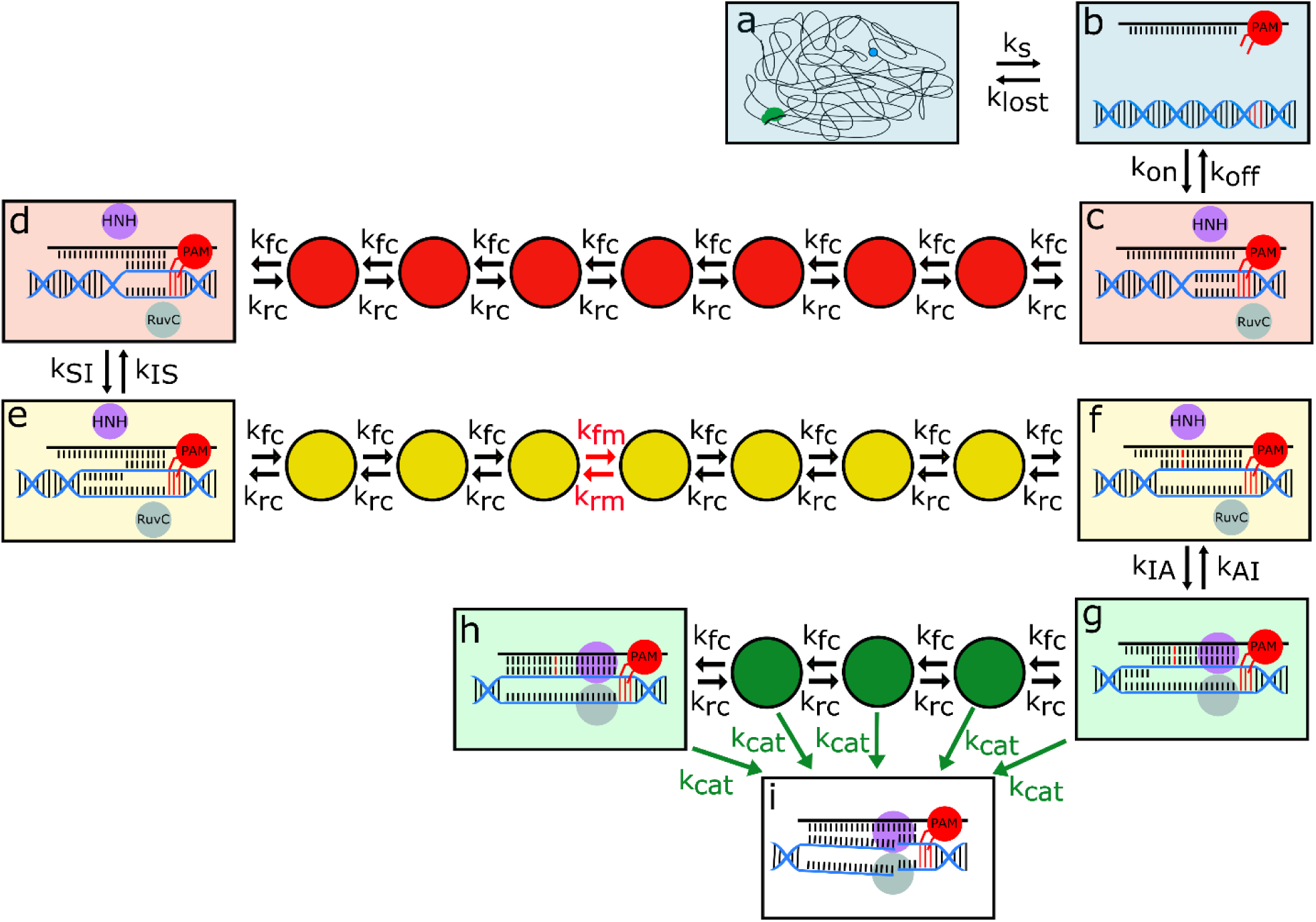
Diagram of the CRISPR-Cas9 target interrogation dynamical system. **(a)** The system starts with CRISPR-Cas9 in the general experimental space, searching for the target. The search rate is the rate of entering the region where the target is located and is defined as k_search_. **(b)** When in proximity with the target, CRISPR-Cas9 can either exit the area of proximity at a rate of k_lost_ or bind to the target PAM with a rate of k_on_. **(c)** Upon forming a PAM bond, CRISPR-Cas9 can initiate the strand displacement of ‘seed’ dsDNA at a rate of k_fc_ for a complementary bond and k_fm_ for a non-complementary bond. The reversal rates for a complementary bond and a non-complementary bond are k_rc_ and k_rm_ respectively. **(d)** Upon forming all the ‘seed’ RNA:DNA bonds the CRISPR-Cas9 system is able to unwind the intermediate segment of the target dsDNA at a rate of k_SI_. **(e)** Winding back the intermediate segment of the target dsDNA occurs at the rate k_IS_ and only when zero intermediate RNA:DNA bonds are formed. Again, upon unwinding the system can form new RNA:DNA bonds at the rates k_fc_ and k_fm_ depending on the complementarity of the nucleotides. The diagram presents an example when the 12^th^ RNA:DNA bond is not complementarity and thus the formation of the 12^th^ RNA:DNA occurs at a rate of k_fm_ instead of k_fc_. **(f)** When CRISPR-Cas9 forms all the intermediate bonds, it can unwind the PAM-distal dsDNA and at the same enter the cleavage active conformation. **(g)** In the cleavage active conformation CRISPR-Cas9 can form more RNA:DNA bonds or proceed to cut the target DNA at a rate of k_cat_. **(h)** The fully formed R-loop. **(i)** We approximate that cutting is irreversible and is thus the ultimate state of our model.

Upon forming a PAM bond, the CRISPR-Cas9 system starts displacing the dsDNA DNA:DNA bonds with spacer-target RNA:DNA bonds starting from the PAM-proximal base pairs (Fig 1C). In our model the rate of displacing a single base pair with a matching RNA:DNA bond happens at a rate of *k*_*fc*_, while the reverse process (displacing the RNA:DNA with a DNA:DNA bond) happens at a rate of *k*_*rc*_. Similarly, rate to form an RNA:DNA mismatch is *k*_*fm*_, and the rate to reverse the RNA:DNA mismatch is *k*_*rm*_. The strand displacement rates are approximated by nucleic acid energetics:

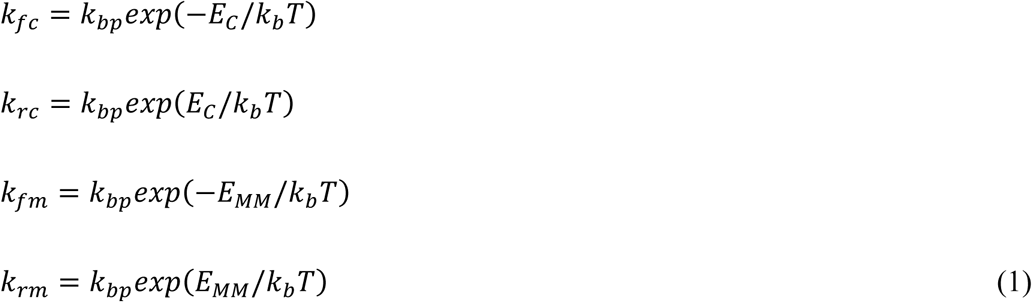

Where *k*_*bp*_ is a normalizing term to set the rate for nucleic acid dynamics, *E*_*C*_ is the average free energy acquired for a matched displacement, and *E*_*MM*_ is the free energy acquired for a mismatched displacement. During the CRISPR-Cas9 strand displacement, the limiting rate is the melting of dsDNA. Experiments studying the dynamics of base pairing approximate the timescales of melting a single DNA:DNA is on the order of 3ms [30] and thus we approximate *k*_*bp*_ as equal to 300 s^-1^. The average energies acquired or lost when replacing a DNA:DNA bond with an RNA:DNA match or an RNA:DNA mismatch are calculated by comparing the energies of DNA:DNA bonds with the energies of RNA:DNA bonds. The average match displacement energy *E*_*C*_ is on the order of 0.5k_b_T, while the average mismatch displacement energy is on the scale of 5k_b_T.

However, the formation of the spacer-target hybrid cannot be modeled by only considering the nucleic acid dynamics. Structural [8,12], smFRET [14,17], and rotor bead tracking experiments [21] demonstrated that during target recognition the Cas9 enzyme undergoes conformational changes that are coordinated by the number of RNA:DNA bonds formed. Therefore, our model accounts for two major conformational changes: the seed to intermediate transition (Fig 1D to Fig 1E) and the intermediate to active conformational change (Fig 1F to Fig 1G). These transitions can only occur when a sufficient number of RNA:DNA bonds are formed and the more of the target DNA can be unwound. The number needed for a seed-intermediate conformational change is *L*_*S*_ and the number of RNA:DNA bonds needed for an intermediate-active conformational change is *L*_*I*_. We set the lengths for *L*_*S*_ and *L*_*I*_ to 10bp and 18bp based on the unwinding of DNA in the magnetic bead study [21] and the rates of the transitions *k*_*SI*_, *k*_*IS*_, *k*_*IA*_, *k*_*AI*_ are all approximated at 1 s^-1^ based on the rates of FRET experiments [17]. However, the rates of the conformational changes could also be on the order of tens of seconds based on results from AFM experiments [31]. CRISPR-Cas9 undergoes conformational changes to enter its catalytically active conformation in which it can cleave the target DNA. In the model the cleavage can happen from any of the states within the active conformation (between Fig 1G and Fig 1H) at a rate of *k*_*cat*_. From kinetic experiments of CRISPR-Cas9, *k*_*cat*_ is on the order of 1 s^-1^ but is highly dependent on the Mg^+2^ of the experimental system [32]. Once the DNA is cut, the system enters the final and irreversible product state (Fig 1I).

From previous CRISPR-Cas9 cleavage kinetics experiments we know that probability of cutting target DNA in CRISPR-Cas9 experiments is approximated with the following formula:

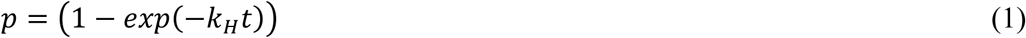

Where *k*_*H*_ is the global cutting rate in a given CRISPR experiment. We can calculate *k*_*H*_ from our model by converting the system of ODEs into a Markov chain (difference equation). We convert the system by assuming that in some timeframe *t*_*s*_ that we set to 1us, the probabilities of forward transitions, *p*_i_, and reverse transitions, *q*_*i*_, can be approximated as:

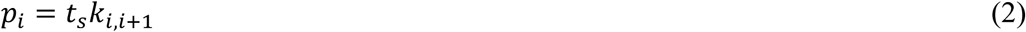

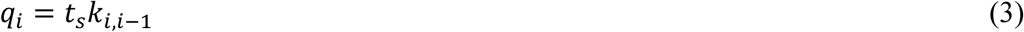

To find the global cleavage rate by taking the inverse of the mean absorption time *τ*_*H*_. Markovian absorption time, *μ*_*0*_, can be found by solving the following system of linear equations:

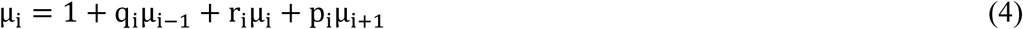

and then scaled to real time using t_s_ as follows:

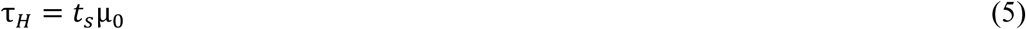

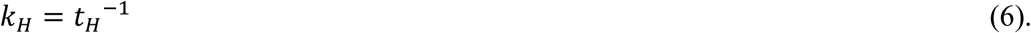

To demonstrate with most clarity the effect conformational changes and target search carry on the mismatch tolerance of CRISPR-Cas9, we calculate the *k*_*H*_ for targets with a single mismatch in each of the 20 possible RNA:DNA bonds (assuming the spacer is 20bp long).

**Table 1.**
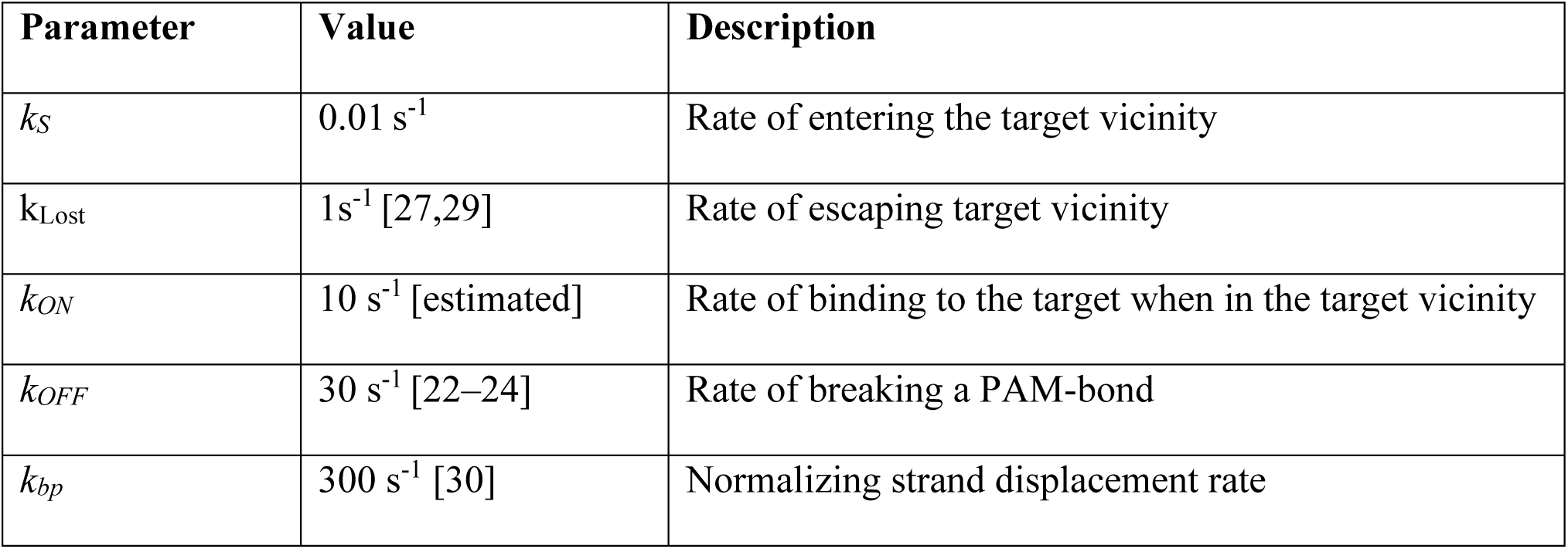
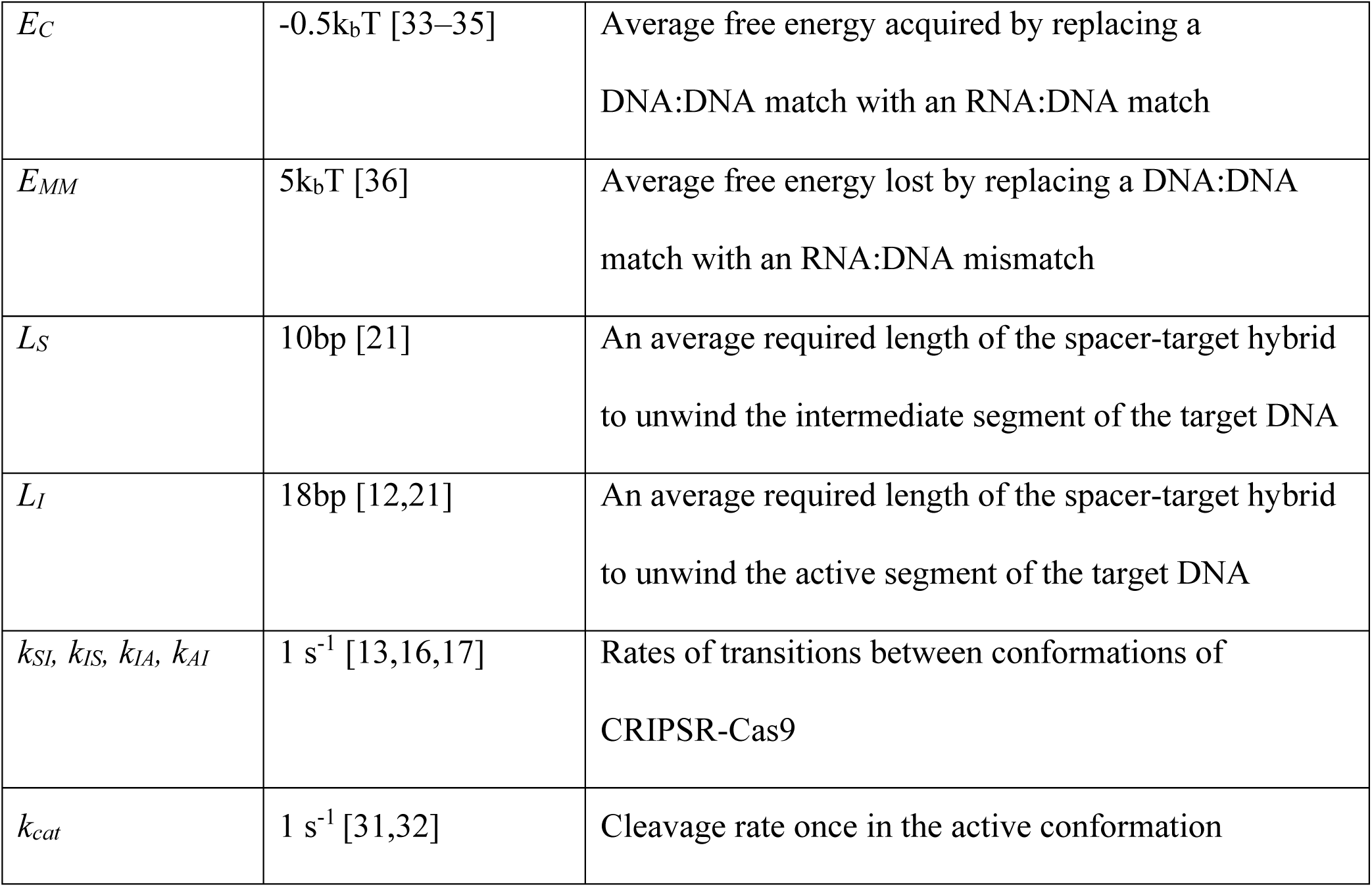
The rates of the CRISPR-Cas9 target interrogation model.

## Results and discussion

### Nucleic acid dynamics during CRISPR-Cas9 target interrogation

Since different target sequences might have different average match and mismatch energies, we observed the effect varying the nucleic acid free energies E_C_ and E_MM_ caried on the mismatch patterns of the model. Predictably, the model demonstrates that decreasing the average free energy acquired when displacing a single dsDNA bond with a matching RNA:DNA bond causes the system to be more tolerant of mismatches (Fig 2A). This is due to the increased stability of the partial RNA:DNA hybrids prior to the mismatch locations, which in turn increase the overall likelihood of surpassing the mismatched site. Similarly, decreasing the energetic penalty E_MM_ increases the likelihood of tolerating a mismatch (Fig 2B), because the relative rate of passing through a mismatch is accelerate in comparison to the rate of unbinding from the DNA. Surprisingly, in this model, the mismatch tolerance patterns are neither uniform, as might be expected for a naïve thermodynamic model, nor are the patterns monotonically increasing as might be expected for a strand displacement kinetic model [25]. To test whether this effect is caused by picking a specific value of k_bp_, we varied the rates from slow to fast (Fig 2C). While changing the rates of k_bp_ affects the mismatch tolerance of the system, the overall pattern remains the same. Another set of nucleic acid parameter are the average conformational change (or unwinding) lengths, which are hypothesized to depend on the sequence of the target DNA [21]. By varying the lengths *L*_*S*_ and *L*_*I*_ in the model, we could control how critical a certain RNA:DNA bond becomes in the target recognition process. When increasing *L*_*S*_ by *1nt* the shape location of the maxima of both mid-spacer peaks also increases by *1nt*. (Fig 2D). However, when increasing the value of *L*_*I*_ by *1nt* only the shape of the second peak changes (Fig 2E).

**Figure 2.**
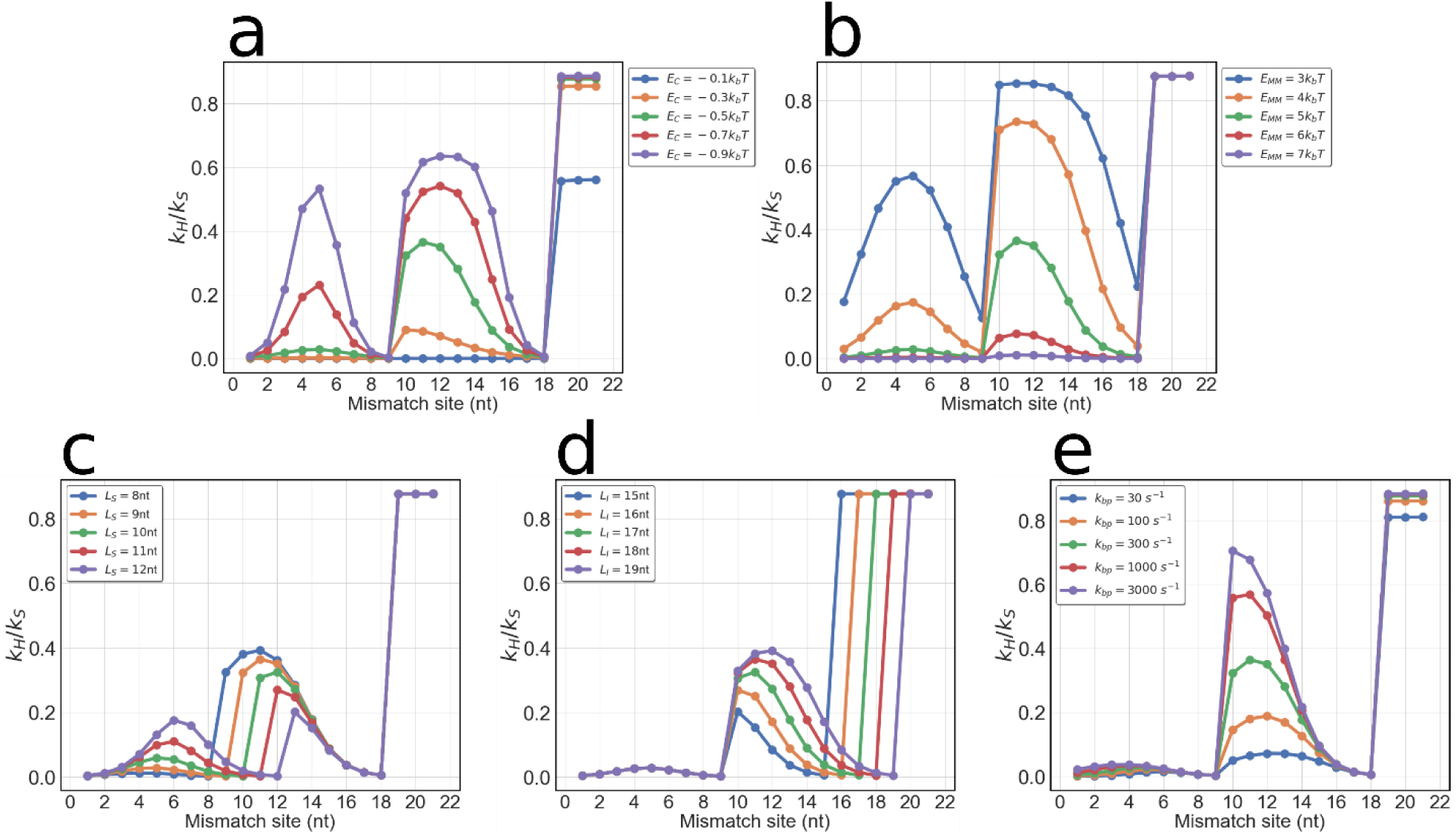
Emergence of single mismatch tolerance in CRISPR-Cas9 systems. The mismatch tolerance patterns of mechanistically faithful CRISPR-Cas9 model are not a monotonic, but instead have multiple local maxima points dependent on the conformational change lengths. **(a)** The energetic parameter E_C_ corresponds to the average free energy acquired for each DNA:DNA bond replaced with an RNA:DNA bond. Decreasing E_C_ makes the system more stable per bond, which in turn increases the likelihood of tolerating a mismatch. **(b)** The other energetic parameter, E_MM_, corresponds to the energetic penalty of a single DNA:DNA matching bond replaced with an RNA:DNA mismatch. As the penalty energy is increased the system becomes more specific, since the energy acquired by matches cannot compensate for the mismatch as effectively. **(c)** The nucleic acid basal rate parameter, k_bp_, plays an important role in the mismatch patterns of CRISPR-Cas9 binding. As we increase k_bp_ the mismatch tolerance also increases due to 2 reasons: (i) a larger k_bp_ increases the relative probability of forming an initial RNA:DNA bond instead of unbinding from the target DNA and (ii) the speed of overcoming a mismatch, k_fm_, becomes relatively large in comparison to the rates of conformational changes and thus are ignored during target recognition. **(d) and (e)** demonstrate the effect changing the conformational change lengths L_S_ and L_I_ respectively have on the single mismatch profiles of CRISPR-Cas9. We notice that mismatches after conformational changes are tolerated the least, since those cause the probability density of the system to be concentrated RNA:DNA hybrid lengths in which the system can revert to the previous conformation.

### Target search and re-binding of CRISPR-Cas9

Normally when modeling the search of a DNA binding enzyme, we approximate the rate of finding the target by some constant rate, k_search_. Upon unbinding from the DNA, the enzyme returns to its unbound state and will re-bind to the target DNA with the same rate k_search_. However, this approach does not take into consideration that when an enzyme re-binds to the target, the rate must be accelerated since the initial location is proximal to the target itself [28]. In the case of Cas9, this effect is amplified by the fact that target search for CRISPR-Cas9 is divided into 1-D diffusion between non-target PAM sites and 3-D diffusion in the cell/tube/microfluidic device (Fig 3A) [23,27]. If Cas9 unbinds from its target PAM site, it is likely to return to the starting PAM site much faster than would be proposed by k_search_. For this reason, in the model we split the target search and binding process into two steps: (i) searching for the general area of the target DNA and (ii) binding to the target DNA when in that region.

**Figure 3.**
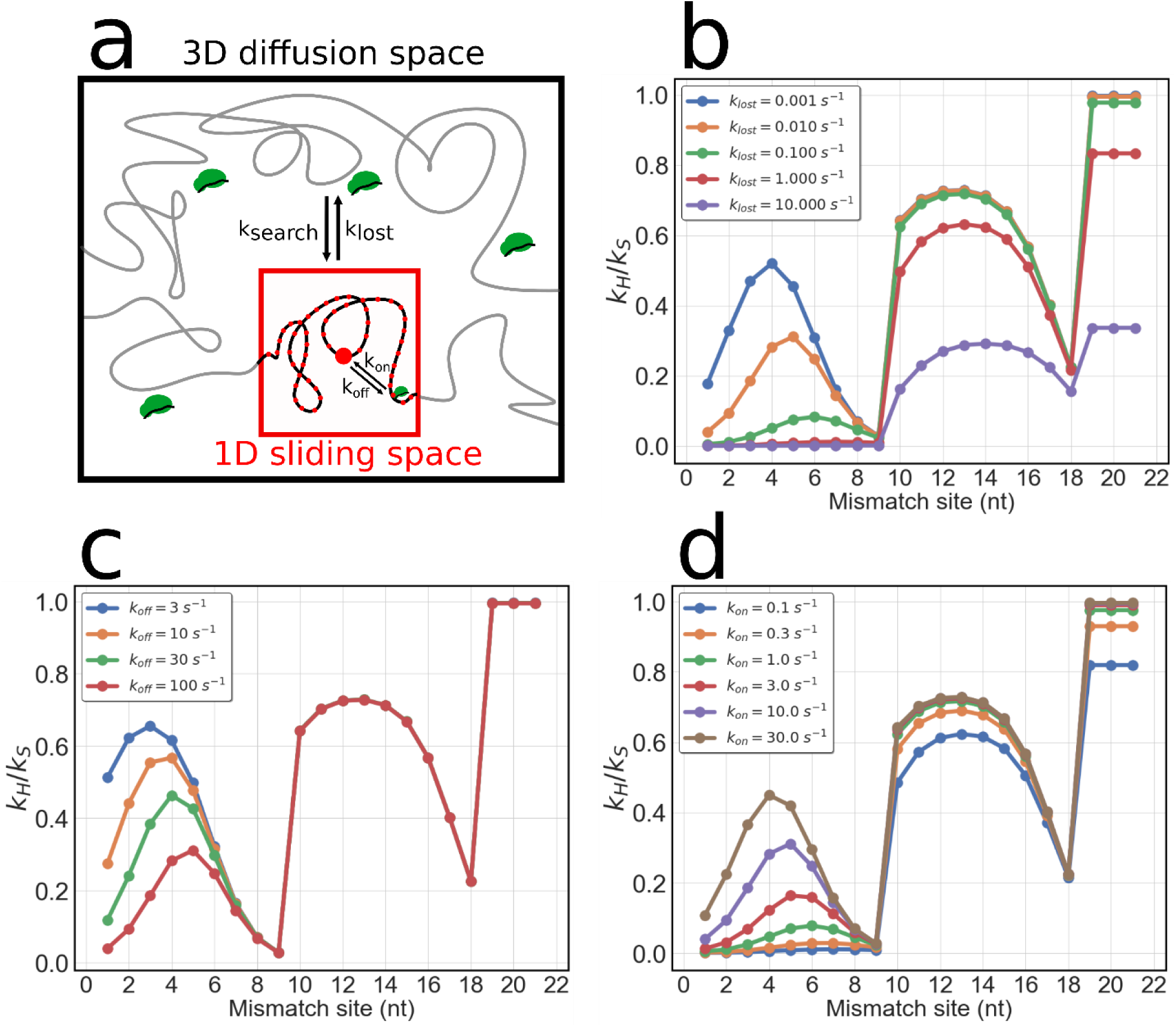
Separation of CRISPR-Cas9 target search in 2 steps. **(a)** A schematic of the difference between 3D diffusion space and the 1D sliding space during CRISPR-Cas9 target search. The search for the target-proximal region occurs with a rate of k_search_, while escaping the target-proximal region occurs with a rate of k_lost_. Within the target proximal region, the motion of CRISPR-Cas9 can be approximated as 1-D sliding between PAM sites. The rate of rebinding to the target within the region is approximated with a single on-rate k_on_ and the unbinding from the target site occurs with some rate k_off_ defined by the energetics of the bond. **(b)** Decreasing k_lost_, causes not only an emergence of PAM-proximal mismatch tolerance, but an increase in the mismatch tolerance as a whole. **(c)** Decreasing k_off_ causes the system to be more tolerance of PAM-proximal mismatches but does not significantly affect the rates of the targets with mismatches past L_S_. **(d)** Increasing the rate k_on_ strongly decreases the specificity of the PAM-proximal bonds, while decreasing the specificity of the mid-target and PAM-distal bond to a lesser extent.

We then tested the effect the diffusion and binding parameters had on the global cleavage rates from equation (6) for single mismatch experiments. Depending on the site, the rate of escaping the proximity of the target might vary. To see how the escape rate affects the single mismatch patterns in the model we varied the rate *k*_*lost*_ (Fig 3B). We notice that *k*_*lost*_ mostly affects the mismatch tolerances of the PAM-proximal RNA:DNA bonds, showing that for smaller rates of 1-D escape, the tolerance for mismatches increases. We then varied the rate of unbinding from a target site can *k*_*off*_ and noticed that it again affects only the PAM-proximal binding to a significant level, while the middle and PAM-distal RNA:DNA bonds appear to maintain the same mismatch tolerance (Fig 3C). We noticed that in a system with a quicker 1-D rebinding rate, *k*_*on*_ the mismatch tolerance also increases only for the PAM-proximal RNA:DNA bonds (Fig 3D).

### Role of DNA unwinding rates in CRISPR-Cas9 target interrogation

When we increase the seed-intermediate transition rate, k_SI_, we observe that the system becomes overall more error prone in comparison to the fully matched target (Fig 4A). Increasing the rate of the reverse process, *k*_*IS*_, slows down the interrogation process and thus decreases the rate of cutting for mismatched targets, without significantly affecting the cleavage rate of a fully matched target (Fig 4B). For the intermediate-active transition, increasing k_IA_ also reduced the specificity of the system but only for the base pairs in the region between L_S_ and L_I_ (Fig 4C). Varying the rate of the reverse process, k_AI_, did not affect the mismatch profile significantly (Fig 4D). This is because the rate of forming the base-pairs after L_I_ is faster than k_AI_ and thus the fully formed target-spacer hybrid serves as a sink in the dynamical system. Unsurprisingly, when increasing the cleavage rate k_Cat_ the model demonstrates that CRISPR-Cas9 becomes less specific (Fig 4E) as demonstrated previously in experiments with the enhanced specificity mutants of Cas9.

**Figure 4.**
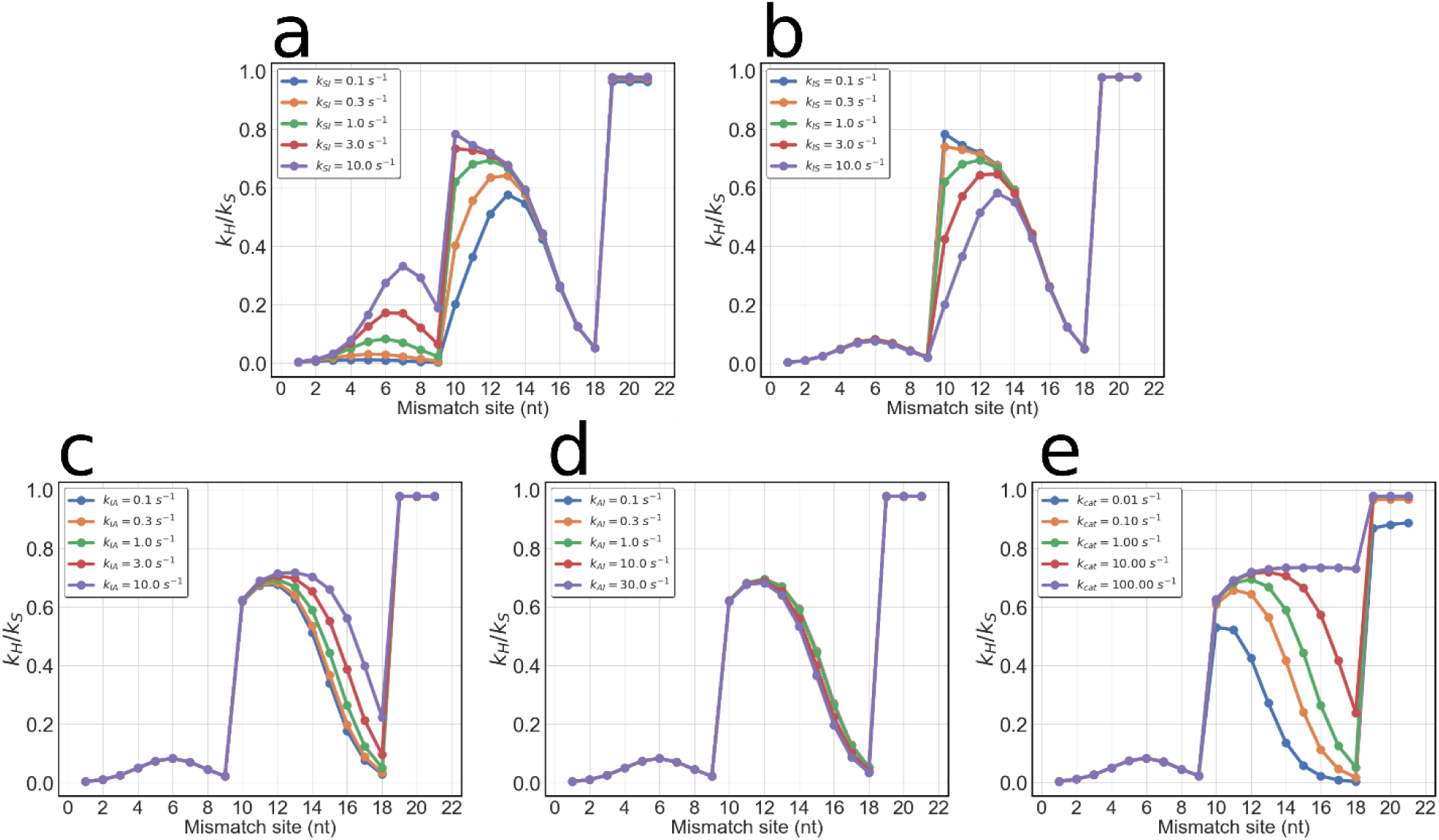
Rates of CRISPR-Cas9 conformational changes/unwinding on mismatch patterns. **(a), (b), (c), and (d)** are the mismatch profiles produced by the model when varying the rates k_SI_, k_IS_, k_IA_, and k_AI_, respectively. **(e)** The mismatch profiles produced by the model for different rates of k_cat_.

We can notice that as we increase the ‘gatekeeping’ rates, k_SI_ and k_IA_, the system reaches the product state faster, but also loses specificity. The mechanism of CRISPR-Cas9 leverages a speed-accuracy tradeoff: the slower the conformational changes, the larger the expected number times a mismatched nucleotide is interrogated. With current directed evolution and rational protein design techniques it might be possible to re-engineer Cas9 (or other Cas enzymes) to have slower or faster transition rates. Therefore, it might be of interest for future work to derive the optimal transition rates as a function of the CRISPR enzymes concentration, spatially dependent diffusion coefficient, and sequences of off-target sites.

## 4 Conclusion

Our model demonstrates that it is unnecessary to assign energetic costs to each separate mismatch of a target to explain the patterns seen in CRISPR-Cas9 off-target experiments. The mismatch patterns can be explained as an emergent effect of the dynamics and topology of the system. However, this model uses only approximate rates and models the target recognition process of CRISPR-Cas9 without considering the spatial effects of the Cas9 enzyme and the nucleic acids. Thus, our model only serves as a semi-quantitative picture of the CRISPR-Cas9 target interrogation to convince CRISPR-Cas9 community to incorporate the dynamics and mechanism of the Cas9 enzyme into off-target prediction software [37].

